# Structural insights into YheS-mediated release of SecM-arrested ribosome

**DOI:** 10.1101/2025.08.28.672766

**Authors:** Kaishi Iso, Toma Ikeda, Kohei Yamasaki, Yushin Ando, Fumiya K. Sano, Tadaomi Furuta, Hideki Taguchi, Osamu Nureki, Yuhei Chadani, Yuzuru Itoh

## Abstract

ATPase-binding cassette subfamily F (ABCF) proteins interact with the ribosome to resolve translation defects near the peptidyl-transferase center, induced by ribosome-targeting antibiotics or “hard-to-translate” nascent peptides. In *Escherichia coli*, four ABCF proteins resolve the translation of distinct problematic sequences, but their molecular mechanisms remain unclear. Here, we present a 2.8-Å cryo-EM structure of the SecM-stalled ribosome in complex with an ATPase-deficient mutant of YheS, an ABCF protein that releases ribosomes stalled during the translation of SecM, a representative ribosome-arresting peptide. YheS is bound to the E-site and relocates the P-site tRNA (P-tRNA) by displacing its CCA-end. Notably, the density of the SecM nascent chain mostly disappeared upon YheS binding. Molecular dynamics simulations support that the P-tRNA relocation generates a pulling force that disrupts key intratunnel interactions essential for arrest. This mechanism is functionally analogous to, yet mechanistically distinct from, the translocon-mediated external pulling force for SecM release.

**Teaser:** The release mechanism of the SecM-arrested ribosome by the ABCF protein YheS was elucidated by structure-based analyses.

## Introduction

The immense diversity of amino acid sequences underpins the functional complexity of proteins. However, it has become evident that ribosomes have difficulties with polymerizing certain amino acid sequences, including stretches of proline, tryptophan, charged residues, or ribosome-arresting peptides, due to interactions within the peptide exit tunnel (**1–15**). Genome-wide and structural studies suggest such problematic sequences are widespread and become more deleterious under stress or aging conditions (**16-21**).

Remarkably, cells have evolved to harness such difficult events for gene regulation. In *E. coli*, the SecM nascent-polypeptide chain arrests the ribosome through interactions within the exit tunnel, thereby promoting the expression of the downstream translocase SecA (**11, 22–27**). The stalled ribosome is subsequently released by an external pulling force generated by the Sec translocon, exemplifying a mechanical strategy for programmed translation arrest.

To minimize the risks posed by such problematic sequences, cells have also evolved specialized translation factors. One example is EF-P (or eIF5A in eukaryotes), which alleviates stalling at polyproline stretches by binding to the ribosomal E-site (**4, 5, 28**). In parallel, members of the ABCF (ATP-binding cassette subfamily F) family have emerged as additional modulators of translation difficulties (**29, 30**). Among them, antibiotic resistance (ARE)-ABCFs dislodge antibiotics from the exit tunnel by inserting a long interdomain linker (∼110 aa) that relocates the P-site tRNA (**31-35**). In contrast, non-ARE ABCFs, comprising EttA, Uup, YbiT, and YheS in *E. coli*, possess shorter interdomain linkers (∼80 aa), termed the P-site tRNA Interacting Motif (PtIM). While EttA promotes elongation immediately after initiation (**36-38**), the functions of the other non-ARE ABCFs have remained unclear.

Recently, we and others have demonstrated that these four ABCFs manage the “hard-to-translate” amino acid sequences (**39-42**). A key finding is that the four ABCFs exhibit “sequence specificity”, targeting distinct challenging motifs. For example, YheS, but not the other ABCFs, specifically releases the SecM-arrested ribosome. Despite their predicted structural similarity, the molecular basis for the YheS-specific release of SecM-induced translation arrest has remained elusive.

In this study, we investigated the mechanism by which YheS alleviates SecM-induced ribosome stalling, using single-particle cryogenic-electron microscopy (cryo-EM) and molecular dynamics (MD) simulations. We determined the structure of the SecM-arrested ribosome in complex with the ATPase-deficient YheS mutant (EQ_2_) at 2.82 Å resolution. YheS occupies the ribosomal E-site and its PtIM protrudes toward the peptidyl transferase center (PTC), forming extensive contacts with the 50S large subunit (LSU) and relocating the P-site tRNA. This induces a ∼3 Å retraction of the SecM nascent chain. Together with MD simulations, our findings suggest that YheS releases the SecM-arrested ribosome through a mechanism distinct from the pulling force generated by the Sec translocon.

## Results

### Overall structure of YheS bound to the SecM-arrested ribosome complex

ABCF proteins act on the ribosome to resolve translational defects, and subsequently hydrolyze ATP upon their dissociation (**29, 36**). To prevent dissociation from the ribosome, we used an ATPase-deficient mutant of YheS (E175Q/E456Q, referred to as EQ_2_) (**40**). The SecM-arrested ribosome was reconstituted using the *E. coli* cell-free PURE system (**43**), and then combined with the purified YheS-EQ_2_ mutant. The resulting complex was directly applied to cryo-EM grids without further purification. Cryo-EM particles were classified based on the rotation state of the 30S small subunit (SSU) and the occupancy at the A, P, and E sites, using focused masks targeting these regions (**fig. S1**). We successfully obtained the cryo-EM map of the SecM-arrested ribosome with the A- and P-site tRNAs, mRNA, and YheS, using 50,797 particles, at an overall resolution of 2.82 Å (**Fig. 1, A and B, fig. S2, Table S1**).

**Fig. 1.**
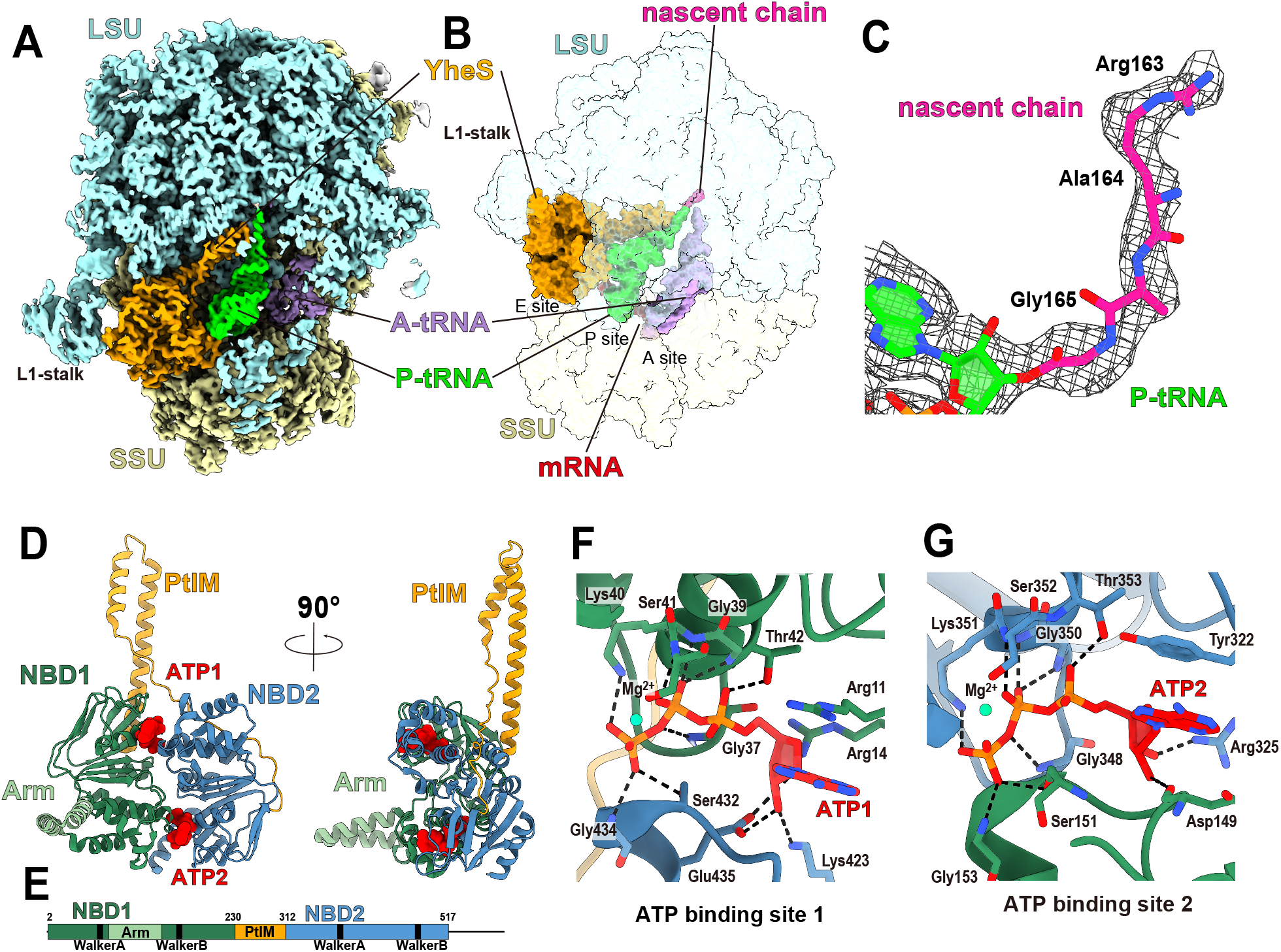
Cryo-EM structure of SecM-arrested ribosome in complex with YheS. (**A**) Overall cryo-EM map of the 70S ribosome with a cutaway view highlighting bound YheS. A- and P-tRNAs are present, and YheS is in the E-site. (**B**) Surface rendering of the complex, showing YheS, A- and P-site tRNAs, mRNA, and the SecM nascent chain. The 70S ribosome is displayed as a transparent surface. (**C**) Close-up view of the nascent SecM chain attached to the P-site tRNA. Only the C-terminal three residues (Arg163–Ala164–Gly165) are resolved. Cryo-EM density is shown as a mesh. (**D, E**) Overall structure and domain composition of YheS. The two nucleotide-binding domains (NBD1 and NBD2) are connected by the P-site tRNA Interacting Motif (PtIM). Two bound ATP molecules are shown as spheres. (**F, G**) Detailed views of the ATP-binding sites in YheS. Mg^2+^ ions are coordinated at both sites. ATP molecules are located at the interface between NBD1 (**F**) and NBD2 (**G**).

YheS binds to the ribosomal E-site and interacts with multiple components, including the 23S rRNA near the E-site and the peptidyl transferase center (PTC), the L1-stalk, uS7 of the SSU head, and the P-site tRNA (P-tRNA). Within the exit tunnel, only the C-terminal three residues of the SecM nascent chain (Arg163-Ala164-Gly165) were resolved, while upstream segments were disordered (**Fig. 1C**). Structurally, YheS forms a pseudo-dimer composed of two nucleotide-binding domains (NBD1 and NBD2), each containing conserved ABC motifs, including the Walker A (P-loop), Walker B, H-switch, Q-loop, D-loop, A-loop, and signature motifs (**Fig. 1, D and E**) (**44**). NBD1 also contains an “Arm” subdomain (residues 90–121) that interacts with the L1-stalk. The two NBDs are connected by a long helix-turn-helix element, PtIM (residues 230–312), which protrudes toward the PTC and interacts with both the 23S rRNA and the acceptor stem of the P-tRNA. The C-terminal extension of YheS (residues 527–637) is disordered.

### ATP interactions in YheS

Two ATP molecules (ATP1 and ATP2) are bound to the P-loops of NBD1 and NBD2, respectively (**Fig. 1, F and G**). At the ATP1 site, the main-chain amide groups of Gly37, Gly39, Lys40, and Ser41, along with the side chains of Lys40 and Thr42, form hydrogen bonds with the phosphate groups of the ATP molecule (**Fig. 1F**). A Mg^2+^ ion is coordinated by the β and γ phosphates of ATP1, as well as by the side chains of Ser41 and Gln71, further stabilizing the nucleotide binding. Additionally, the side chain of Arg14 interacts with the ribose moiety of ATP1 via a hydrogen bond. ATP2 is coordinated similarly by the corresponding conserved residues. While both ATP sites share a conserved binding mode, there are notable differences: Arg11 forms a π-stacking interaction with the adenine of ATP1, whereas the corresponding interaction at the ATP2 site is mediated by Tyr322 (**Fig. 1G**). Furthermore, the side chains of Lys423 and Glu435 form hydrogen bonds with the ribose of ATP1, whereas no such interactions are observed at the ATP2 site.

Importantly, the ATP binding sites are located at the interface between NBD1 and NBD2 (**Fig. 1, D, F and G**). The γ-phosphate of ATP1 forms additional hydrogen bonds with the side chain of Ser432 and the main chain of Gly434 from NBD2; equivalent interactions are mirrored at the ATP2 site with the corresponding residues in NBD1. An additional hydrogen bond is formed between Asp149 of NBD1 and the ribose of ATP2. Because these interdomain interactions are mediated by the γ-phosphate, ATP hydrolysis is expected to disrupt the NBD1–NBD2 interface. This would induce a conformational change of YheS, likely leading to its dissociation from the ribosome.

### Arm interactions with the L1-stalk of LSU

The Arm subdomain of YheS interacts with both the protein and RNA components of the ribosomal L1-stalk of LSU, specifically uL1 and helix 76 (H76) of the 23S rRNA (**Fig. 2, A and B, Table S2**). Residues His114, Asp118, and Asp121 form hydrogen bonds or salt bridges with uL1, while Asn105 and Trp123 of YheS form stacking interactions with the uL1 residues. In parallel, Ser127, His134, and Asn140 hydrogen bond with the nucleotides in H76. Although the L1-stalk is typically flexible and thus often poorly resolved in structural studies (**27, 45**), it is well defined in our cryo-EM map (**Fig. 1A**), likely due to the stabilization by these extensive YheS-ribosome interactions. In addition, YheS interacts with helix 68 (H68) of domain IV in the 23S rRNA, adjacent to the L1-stalk. Residues Asn25 and Thr228 in NBD1, along with Gln244 (near the base of PtIM), form hydrogen bonds with the H68 nucleotides (**Fig. 2C**).

**Fig. 2.**
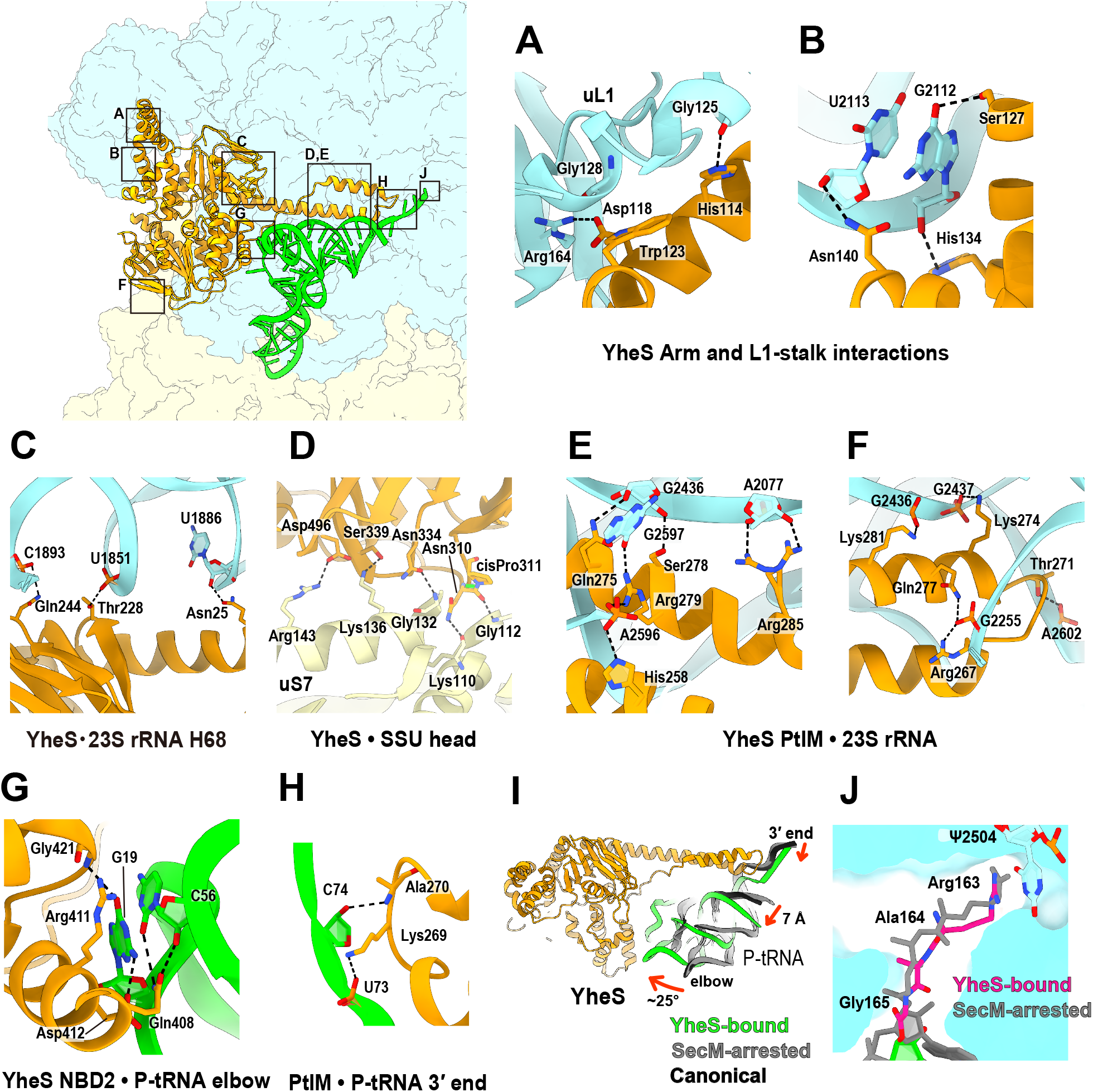
YheS interactions with the ribosome and P-site tRNA. (**A, B**) The Arm subdomain of YheS interacts with both the protein (uL1, **A**) and RNA (helix H76 of 23S rRNA, **B**) components of the L1-stalk. (**C**) NBD1 of YheS interacts with ribosomal protein uS7 in the SSU head. (**D**) The residues at the base of PtIM form multiple polar interactions with the 23S rRNA near the L1-stalk. (**E, F**) PtIM of YheS forms multiple polar interactions with the 23S rRNA near the PTC. (**G, H**) Interactions between YheS and P-tRNA. NBD2 of YheS forms hydrogen bonds and stacking interactions with the P-tRNA elbow (**G**), while the tip of PtIM forms two hydrogen bonds with the 3′ end of the P-tRNA (**H**). (**I**) Relocation of the P-site tRNA upon YheS binding. The P-tRNA in the current complex is colored green, while that of the SecM-arrested ribosome without YheS (PDB: 8QOA) (**27**) and that of the canonical elongating ribosome (PDB: 8B0X) (**45**) are shown in black and green, respectively. (**J**) Conformational comparison of the P-tRNA and SecM nascent chain between the SecM-arrested ribosome with or without YheS (gray; PDB: 8QOA). In the YheS-bound state, the 3′ end of the P-tRNA and the attached SecM nascent chain are retracted by ∼2 Å toward the tunnel entrance.

### Interactions with the SSU head and the LSU central protuberance

NBD1 of YheS interacts with the ribosomal protein uS7 in the head of the SSU (**Fig. 2D and fig. S3**). Residues Asn310, Pro311, Asn334, Ser339, and Asp496 form hydrogen bonds or salt bridges with uS7. These interactions appear to induce a ∼6 Å shift of the SSU head, resulting in expansion of the E-site space compared to the YheS-free ribosome (**fig. S3A**). A cis-peptide bond between Asn310 and Pro311 likely contributes to the formation of this specific interface (**Fig. 2D**). YheS also interacts with the central protuberance (CP) of the LSU, displacing it by ∼5 Å (**fig. S3A**). Although no obvious hydrogen bonds are observed, NBD2 of YheS and the ribosomal protein uL5 in the CP are located within the van der Waals interaction distance (**fig. S3B**). Since the CP normally interacts with the elbow of the P-tRNA (**fig. S3B**), this CP shift is likely related to the P-tRNA relocation induced by YheS, as described below. These shifts in the SSU head and the LSU CP eliminate their inter-subunit contacts, particularly those between uS13 (SSU) and uL5/bL31 (LSU), resulting in a spatial gap in this region (**fig. S3A**).

### PtIM interactions with the 23S rRNA

YheS establishes extensive contacts with domain V of the 23S rRNA near the PTC (**Fig. 2, E and F, Table S2**). Specifically, ten residues located at the tip of PtIM (His258, Arg267, Thr271, Lys274, Gln275, Gln277, Ser278, Arg279, Lys281, and Arg285) form hydrogen bonds or salt bridges with the nucleotides in helices 74, 75, 80, and 93 of domain V. These interactions anchor PtIM firmly to the LSU and define its position relative to the PTC. A comparison with the SecM-arrested 70S ribosome in the absence of YheS (PDB ID: 8QOA) (**27**) reveals that the overall conformation of the 23S rRNA near the PTC remains largely unchanged, except for several tunnel-lining nucleotides (A2062, U2506, U2585) (**fig. S3C**), as well as A2602 in helix H93 (**fig. S3D**). Specifically, A2062 and U2506 change their base orientations, U2585 moves outward from the exit tunnel, and A2602 adopts a flipped-in conformation. These movements appear to result from the shifted acceptor stem of the P-tRNA, as described below.

### P-tRNA interaction and relocation

YheS interacts with the P-tRNA at multiple regions, contributing to its relocation. In NBD2, Gln408 forms hydrogen bonds with nucleotide C56 of the P-tRNA, and Arg411 stacks onto the base via a π-interaction (**Fig. 2G**). In parallel, Asp412 and Gly421 form hydrogen bonds with G19. YheS also binds to the 3′ region of the P-tRNA. Lys269 and Ala270, located at the tip of PtIM, form a salt bridge with U73 and a hydrogen bond with C74, respectively (**Fig. 2H, Table S3**).

Compared to the SecM-stalled 70S ribosome lacking YheS (PDB: 8QOA), the position of the P-site tRNA acceptor stem is drastically altered (**Fig. 2I**). The tip of PtIM occupies the canonical location of nucleotides 71–74 in the acceptor stem. Notably, the position of PtIM is stabilized by multiple interactions with the 23S rRNA (**Fig. 2, E and F**). As a result, the P-tRNA acceptor stem is displaced by ∼7 Å, disrupting the base pair between C75 of the P-tRNA and G2251 of the 23S rRNA (**fig. S3D**). In parallel, the elbow region of the P-tRNA rotates by ∼25° (**Fig. 2I**), likely due to the interaction between YheS and the P-tRNA elbow. This rotation also alters the interaction with the CP: in the SecM-arrested ribosome without YheS, Arg80 of uL5 stacks with the elbow of the P-tRNA, but this interaction is lost in the YheS-bound complex (**fig. S3B**). Instead, Arg80 forms a salt bridge with G57 of the P-tRNA (**fig. S3B**). This change arises from both the P-tRNA rotation and the CP displacement induced by YheS binding.

As consequences of these structural rearrangements, the 3′-terminal A76 of the P-tRNA is shifted outward by ∼3 Å from the PTC (**Fig. 2J**), applying tension to the covalently linked SecM nascent chain. In addition, the density for the SecM nascent chain is weak, and only the C-terminal three residues (Arg163-Ala164-Gly165) can be traced (**Figs. 1C and 2J**). These findings collectively suggest that YheS binding relocates the P-tRNA along with the attached SecM nascent chain, thereby disrupting the intratunnel interactions required for translation arrest (**27**).

### Mutational analysis of YheS–ribosome and tRNA interactions

To evaluate the functional significance of the YheS interactions with the ribosome and P-site tRNA, we introduced a series of point mutations into YheS and measured their ability to release SecM-arrested ribosomes (**Fig. 3**). Notably, the single-point substitutions D118A and W123A, which disrupt the interactions with the ribosomal protein uL1, and R279A, targeting the 23S rRNA near the PTC, each led to a greater than 50% reduction in activity (**Fig. 3, A and F**). The corresponding multi-point mutants, H114A/D118A/W123A and Q275A/Q277A/R279A/K281A, exhibited even more severe defects, supporting the importance of these contact sites. In contrast, mutations targeting YheS interactions with more distal regions of the 23S rRNA or with uS7 in the SSU resulted in minor reductions in activity (**Fig. 3, B, C, and G**). These findings suggest that the direct contacts with the protein components of the L1-stalk and with the 23S rRNA adjacent to the PTC affect the YheS function.

**Fig. 3.**
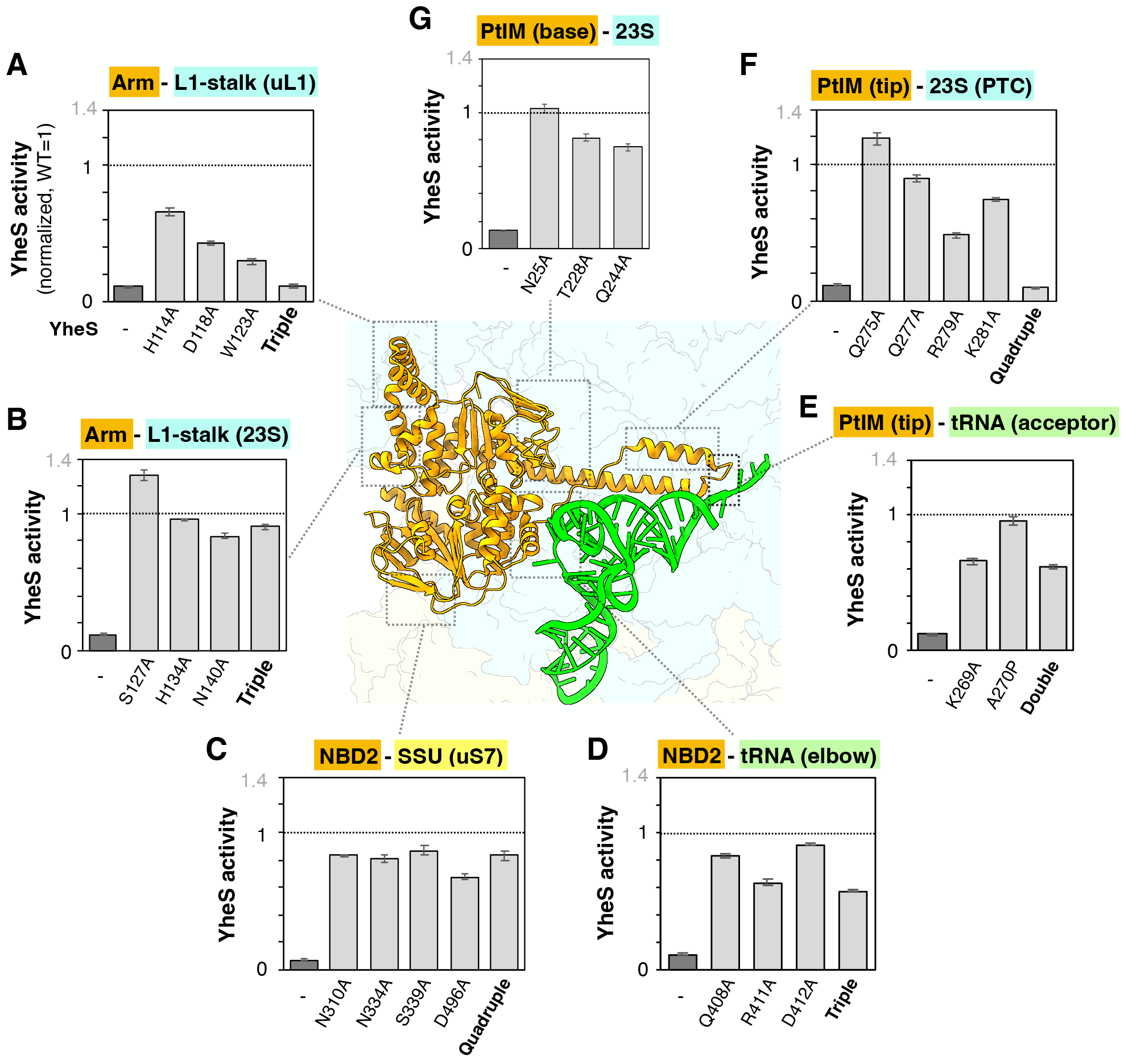
Mutational analysis of YheS residues. Alanine or proline substitutions were introduced into YheS residues that mediate interactions with the ribosome or P-tRNA, either individually or in combination. The arrest-releasing activity of each mutant was quantified as described in the Methods and normalized to wild-type YheS (set to 1). The mean ± SE estimated from three independent biological replicates is shown. (**A**) The Arm-uL1 interaction in the L1-stalk (H114A, D118A, W123A, triple mutant). (**B**) The Arm-rRNA interaction in the L1-stalk (S127A, H134A, N140A, triple mutant). (**C**) NBD1-uS7 interaction in the SSU head (N310A, N334A, S339A, D496A, quadruple mutant). (**D**) NBD2 residues contacting the P-tRNA elbow (Q408A, R411A, D412A, triple mutant). (**E**) The PtIM tip contacting the P-tRNA acceptor stem (K269A, A270P, double mutant). (**F**) The PtIM tip contacting 23S rRNA near the PTC (Q275A, Q277A, R279A, K281A, quadruple mutant). (**G**) The base of PtIM contacting 23S rRNA near the L1-stalk (N25A, T228A, Q244A).

Unexpectedly, mutations disrupting YheS interactions with the P-tRNA, whether targeting the elbow region or the 3′ acceptor stem, had only modest effects on its activity (**Fig. 3, D and E**). This indicates that while YheS interacts with the P-tRNA, these interactions are auxiliary, and the primary mechanism of action relies on the ribosome binding. These results, together with the structural observations, support a model in which YheS binds to the E-site through interactions with uL1 and inserts its PtIM near the PTC, thereby displacing the acceptor stem of the P-tRNA along with the attached SecM nascent chain.

### Behavior of SecM during P-tRNA relocation in molecular simulations

To examine how the YheS-induced relocation of the P-tRNA affects the attached SecM nascent chain, we conducted steered molecular dynamics (SMD) simulations of SecM in the tunnel. Simulations were performed starting with a partial complex, extracted from the previously reported SecM-arrested ribosome (**27**), and the SecM segment (residues 149– 165) was extended by 2, 5, and 10 Å to mimic the pulling effect (**Fig. 4, A-D**). Extensions of 2 or 5 Å resulted in only minor perturbations in the C-terminal residues of SecM (**fig. S4A**). In contrast, a 10-Å extension led to pronounced conformational changes. Specifically, the distances between Ile156 and Ile162, as well as between Arg163 and Ψ2504, increased by ∼2 Å (**Fig. 4, B, C, and E**). Time-course analyses revealed that these disruptions occur sequentially. The Ile156–Ile162 distance begins to increase at 51.2 ns, corresponding to a nascent chain extension of less than 4 Å. The Arg163–Ψ2504 distance then increases at 68.1 ns, when the extension reached ∼6 Å (**Fig. 4, D and E**). This order is consistent across five independent simulations.

**Fig. 4.**
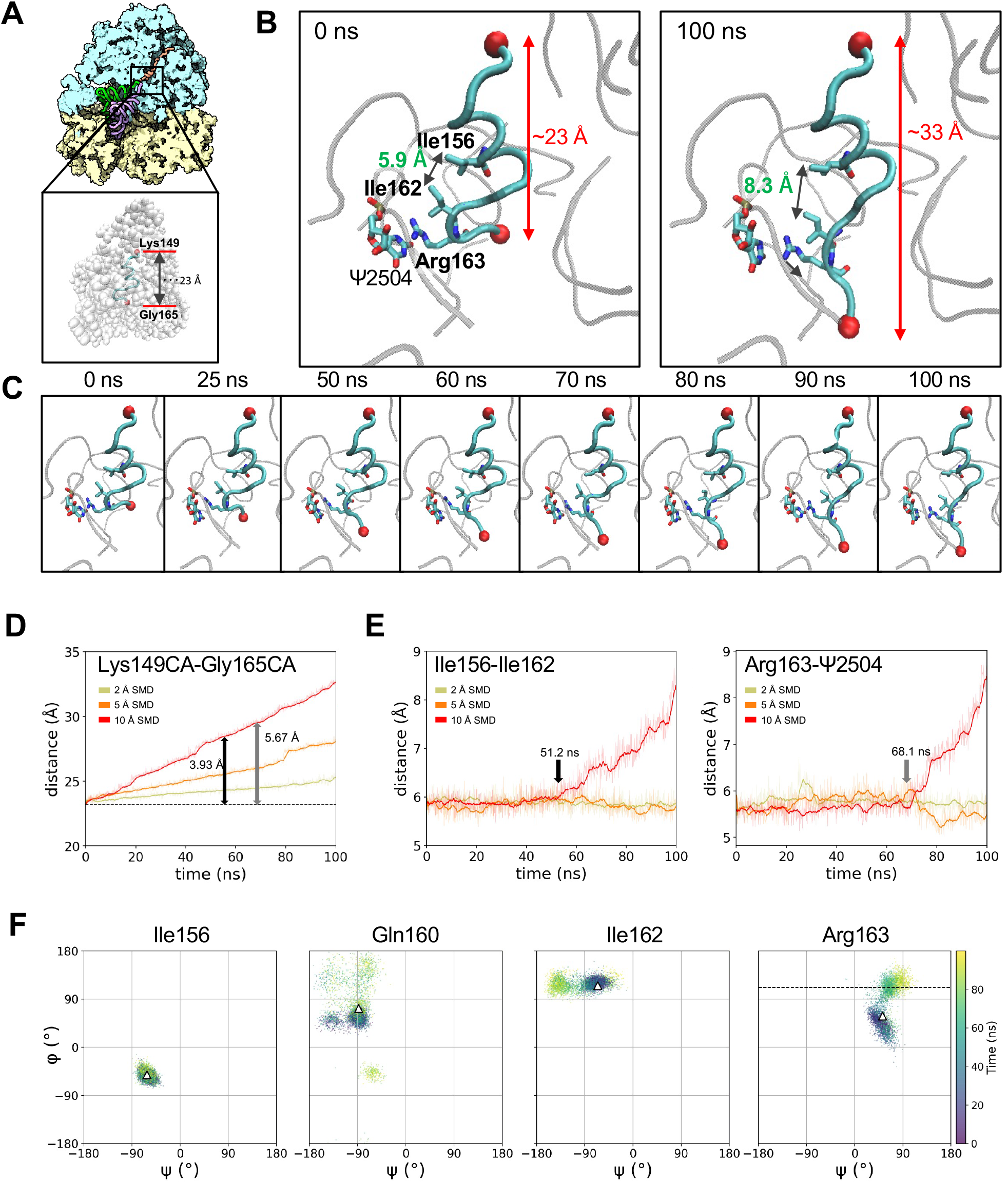
Steered Molecular Dynamics (SMD) simulations of the SecM nascent chain in the stalled ribosome. (**A**) Overall view of the SecM-arrested ribosome complex (PDB: 8QOA, top) and the partial structure used in the simulations (bottom). At the top, the large and small ribosomal subunits (LSU and SSU) are shown as cyan and yellow surfaces, respectively; the P-site tRNA, A-site tRNA, and SecM are shown as green, purple, and orange tubes. At the bottom, the LSU is shown as a white sphere and the modeled SecM segment as a cyan tube. Residues Lys149 and Gly165, used for force application in SMD, are shown as red spheres. (**B, C**) Snapshots of the 10 Å extension simulation. (**B**) Enlarged views at 0 ns and 100 ns. (**C**) Overview at multiple time points during the 100 ns trajectory. Key residues involved in intrachain and ribosome-nascent chain interactions (Ile156, Ile162, Arg163, Ψ2504) are highlighted in sticks. (**D**) Time-dependent changes in the C_α_–C_α_ distance between Lys149 and Gly165 under 2 Å (ochre), 5 Å (orange), and 10 Å (red) extension conditions. The dashed line indicates the initial distance in the starting structure. Black and gray arrows indicate the onset of distance increases between Ile156–Ile162 and Arg163– Ψ2504, respectively, shown in (**E**). (**E**) Time evolution of the Ile156–Ile162 distance (left) and the Arg163–Ψ2504 distance (right) under 2 Å, 5 Å, and 10 Å extensions. (**F**) Changes in backbone dihedral angles of four residues (Ile156, Gln160, Ile162, and Arg163) during five independent SMD simulations with 10 Å extension. Each triangle indicates the initial dihedral angles, and the dashed line in Arg163 represents the dihedral angle observed in the YheS-bound structure.

To evaluate the structural stability of this region, we conducted additional 200-ns conventional MD simulations with positional restraints on the C-terminus of the nascent chain. These simulations highlighted the importance of the hydrophobic interactions between Ile156 and Ile162, as slight increases in structural fluctuations are observed at the neighboring residues Ser157 and Gly161 (**fig. S4B**). Backbone dihedral-angle analyses of the SMD simulations indicated that residues Ser152–Ala159, which constitute the α-helical segment of SecM, remain stable under all extension conditions. In contrast, Gln160, located adjacent to the C-terminus of the helix, undergoes dihedral angle shifts, leading to a partially unfolded conformation in several trajectories (**Fig. 4F and fig. S4C**). In addition, Arg163, which adopts a highly restricted dihedral angle resembling a left-handed helix in the arrested structure, shifts toward a random-coil-like conformation (**Fig. 4F**). The φ angle of Arg163 in the simulations closely matches the corresponding angle observed in the YheS-bound structure in this study.

### Comparison with other ABCF proteins

The key structural difference between Antibiotic Resistant (ARE) ABCFs and non-ARE ABCFs lies in the length of their PtIM (**Fig. 2I, fig. S5**). In the ARE ABCF-ribosome complexes, the long PtIM extends into the peptidyl transferase center (PTC) and even the exit tunnel, displacing antibiotics bound within the tunnel (**fig. S5A**) (**31-35**). Accordingly, ARE-ABCFs with long PtIMs are unable to function when a nascent chain occupies the exit tunnel and are thought to act only when the initiator tRNA is accommodated (**fig. S5A**). In contrast, non-ARE ABCFs such as *E. coli* EttA possess short PtIMs that do not reach the PTC (**37**) (**fig. S5B**). These ABCFs induce only minor rRNA rearrangements or subtle shifts in the P-tRNA, but help stabilize the P-tRNA to promote efficient peptidyl transfer (**38**).

Interestingly, in our YheS-ribosome structure, the P-tRNA is relocated more substantially than in other non-ARE ABCF complexes, resembling the extent seen with ARE-ABCFs (**Fig. 2I, fig. S5**), despite the short PtIM of YheS. Since the SecM nascent chain occupies the exit tunnel, YheS likely induces P-tRNA relocation to generate a pulling force and disrupt the interactions within the tunnel. This is achieved via extensive contacts between PtIM and the 23S rRNA near the PTC (**Fig. 2, E and F**), combined with contacts at the tRNA elbow that rotate the P-tRNA toward the E-site (**Fig. 2G**).

Finally, ATP hydrolysis likely triggers conformational changes at the NBD1– NBD2 interface, disrupting the interdomain interactions mediated by the γ-phosphate groups (**Fig. 1, D–G**). These changes would render YheS incompatible with the E-site architecture and promote its dissociation. Upon YheS release, the P-site tRNA can be relocated correctly, allowing translation to resume its normal cycle (**Fig. 5**).

**Fig. 5.**
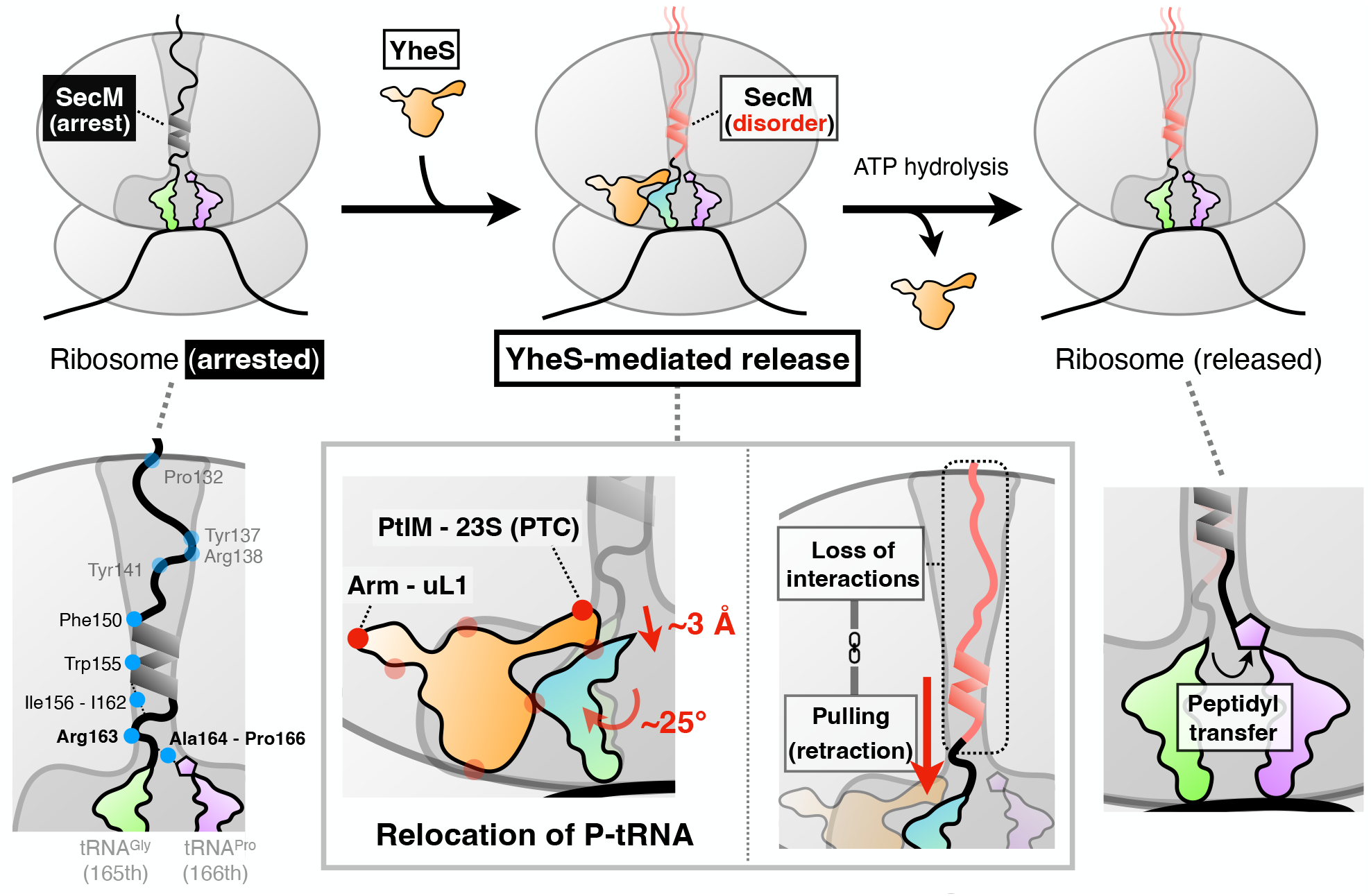
Mechanism of SecM-arrested ribosome release by YheS. Model for the YheS-mediated release of the SecM-arrested ribosome. Upon translation arrest, the SecM nascent chain forms extensive interactions throughout the exit tunnel. YheS binds to the vacant E site, relocates the P-tRNA, and exerts a pulling force on the nascent chain toward the tunnel entrance, disrupting the intratunnel interactions. Subsequent ATP hydrolysis triggers a conformational change of YheS, leading to its dissociation and allowing the nascent chain to adopt a canonical conformation at the PTC.

## Discussion

In this study, we determined the 2.82 Å structure of YheS in complex with the ribosome arrested by the SecM nascent chain. The structure revealed multiple contact points, and the mutational analysis highlighted the importance of the Arm and PtIM of YheS, which interact with the L1-stalk and the 23S rRNA near the PTC. These interactions displace the CCA end of the P-site tRNA by ∼3 Å and retract the SecM nascent chain by ∼2 Å. Importantly, no electron density is observed for residues N-terminal to Arg163, indicating that this region becomes disordered upon YheS binding. In contrast, previous structures obtained in the absence of YheS showed that the SecM nascent chain remains stably folded throughout the exit tunnel (**27**).

A previous study demonstrated that SecM-induced arrest relies on cumulative interactions between the nascent chain and the exit tunnel, including an α-helix downstream of Phe150 and the stacking interaction between Arg163 and Ψ2504 (**27**). These interactions stabilize Ala164 and facilitate hydrogen bonding with the incoming proline at the A-site, thereby inhibiting peptidyl transfer. These residues have been validated as critical for arrest by mutagenesis (**11**). Although Ile156 and Ile162 have also been highlighted, their roles remain unclear due to the absence of direct contacts with the ribosome.

Our MD simulations indicated that YheS binding induces structural rearrangements, including the sequential separation of Ile156–Ile162 and Arg163–Ψ2504, as well as dihedral angle shifts at Gln160 and Arg163. These perturbations likely destabilize additional intratunnel interactions. While the 3′ end of P-tRNA is displaced by only ∼3 Å in the cryo-EM structure, the simulations suggest that larger displacements (∼10 Å) are necessary to fully disrupt the interactions within the tunnel. These findings imply that the observed structure may represent a post-release state formed after the disruption of these critical interactions.

Based on our findings, we propose the following model for YheS-mediated release (**Fig. 5**):

1. Recognition: YheS binds to the vacant E-site via the L1-stalk.
2. Insertion: PtIM of YheS relocates the P-tRNA and forms stable contacts with the 23S rRNA.
3. Disruption: This P-tRNA relocation shifts the attached SecM nascent chain, disrupting key interactions such as Ile156–Ile162 and Arg163–Ψ2504, and thus destabilizing and releasing extensive contacts within the exit tunnel.
4. Dissociation: ATP hydrolysis triggers conformational changes in YheS, promoting its dissociation from the ribosome.

This model may explain why YheS is ineffective against other arrest peptides, such as the RAPP motif in *S. medicae* ApdP (**40**). While ApdP arrests the ribosome through an identical mechanism involving the penultimate alanine and the A-site proline (**46**), it relies more heavily on PTC-proximal residues and less on extended tunnel contacts (**16**). Therefore, YheS, which disrupts interactions within the tunnel, may be ineffective at releasing ApdP-induced arrest. Alternatively, the stalling structure of the RAPP motif may be rapidly reconstructed after YheS dissociates.

While translation factors like EF-P (eIF5A) and EttA promote elongation by stabilizing the P-site tRNA from the E-site (**37, 38, 47**), YheS acts by reshaping the conformation and interactions of the nascent chain within the exit tunnel. This action is also distinct from the ARE-ABCFs, which directly dislodge antibiotics from the tunnel (**31-35**). These findings underscore how different ABCF proteins resolve diverse translational defects via specialized mechanisms (**31-35, 39-42, 48**). Moreover, the actions of YheS contrast with the external pulling force of the Sec translocon, yet both effectively relieve SecM-induced arrest, highlighting the mechanistic versatility in arrest resolution (**26, 48-51**).

Finally, our cryo-EM and MD simulation results provide a mechanistic framework for understanding how YheS resolves SecM-induced stalling. These insights contribute to a broader understanding of the functional specialization of ABCF proteins and may ultimately inform the rational design of synthetic translation regulators tailored to specific sequence or structural contexts.

## Materials and Methods

### *E. coli* strains, plasmids, and oligonucleotides

*E. coli* strain BW25113 {Δ(*araD-araB*)567, Δ*lacZ4787*(::*rrnB*-3), λ-, *rph*-1, Δ(*rhaD*-*rhaB*)568, *hsdR514*} was used as the experimental standard strain. Plasmids and oligonucleotides used in this study are listed in **Tables S4 and S5**, respectively. Plasmids were constructed using standard cloning procedures, including Gibson assembly and site-directed mutagenesis. The sequence files of the plasmids constructed in this study are available in the Mendeley repository [doi: 10.17632/5cjmw3wmp8.1].

### *In vitro* translation

The coupled *in vitro* transcription-translation reaction was performed using PURE*frex* v1.0 (GeneFrontier), as described previously (**10, 52**). The DNA template encoding the SecM arrest peptide was PCR-amplified from pCY3867, using primer 1 (GGCCTAATACGACTCACTATAGGAGAAATCATAAAAAATTTATTTGCTTTGTG AGCGG) and primer 3 (AGTCAGTCACGATGAATTCCCCTAGCTTGG). Following an incubation at 37°C for 30 min, the reaction mixture was treated with purified YheS-EQ_2_ at a final concentration of 5 µM and incubated for an additional 10 min. The mixture was then placed on ice and immediately used for cryo-EM grid preparation.

### Purification of YheS-EQ_2_ protein

*E. coli* BL21 (DE3) cells harboring pCY3863, which encodes N-terminally His_6_-tagged YheS-EQ_2_, were grown overnight at 37°C in LB medium supplemented with 100 µg/ml ampicillin. The next day, the overnight culture was diluted into fresh LB medium containing 100 µg/ml ampicillin and grown at 37°C until the A_600_ reached 0.6. Subsequently, 0.05% arabinose was added to express YheS-EQ_2_ proteins, and the culture was further cultivated for 2 hours at 37°C. Then cells were harvested by centrifugation at 5,000 x g for 15 min at 4°C.

Cell pellets were washed and resuspended in disruption buffer (20 mM Tris-HCl, pH 7.5, 500 mM NaCl, 10% glycerol, 7 mM β-ME, 10 mM imidazole, EDTA-free cOmplete^TM^ mini protease inhibitor cocktail). Cells were disrupted by sonication, and debris was removed by ultracentrifugation (100,000 x g, 40 min at 4°C). The recovered lysate was loaded onto a column filled with Ni-NTA Superflow resin (QIAGEN). After washing the column with ten column volumes of wash buffer {20 mM Tris-HCl, pH 7.5, 500 mM NaCl, 10% glycerol, 7 mM β-ME, 20 mM imidazole}, the YheS-EQ_2_ protein was eluted by wash buffer containing steps of 50, 100, and 300 mM imidazole. The fraction containing the YheS-EQ_2_ protein was concentrated using an Amicon Ultra 30 kDa filter (Merck) and solvent exchanged on a PD-10 column (Cytiva) to storage buffer (20 mM Tris-HCl, pH 7.5, 150 mM NaCl, 10% glycerol, 7 mM β-ME). The purified protein was stored at −80°C.

### Grid preparation and cryo-EM data acquisition

For the cryo-EM analysis, 3 μL of the PURE system reaction solution containing stalled ribosomes was applied onto a glow-discharged holey-carbon grid coated with a continuous 2 nm-thick carbon film (Quantifoil Au 300 mesh, R2/1 +2 nm Ultra-Thin Carbon Film) and incubated for 30 s in a controlled environment of 100% humidity at 4°C. The grids were blotted for 4 s, and then plunge frozen in liquid ethane, using a Vitrobot Mark IV (FEI). The datasets were collected using a Titan Krios G4 microscope (Thermo Fisher Scientific), running at 300 kV and equipped with a Gatan Quantum-LS Energy Filter (GIF). A Gatan K3 Summit direct-electron detector was used at a pixel size of 0.83 Å (magnification of ×105,000) with an exposure of approximately 30 electrons per Å^2^ with 30 movie frames. The data were automatically acquired using the EPU software (Thermo Fisher Scientific), with a defocus range of −0.8 to −2.0 μm, and 18,200 movies were obtained.

### Cryo-EM image processing

The data were processed with cryoSPARC (**53**) and RELION (**54**). In cryoSPARC, the dose-fractionated movies were subjected to beam-induced motion correction and dose weighting using patch motion correction, and the contrast transfer function (CTF) parameters were estimated using patch-based CTF estimation. From the 18,200 preprocessed micrographs, 1,770,974 particles were automatically picked by the template picker, using the references created by the blob picker and several rounds of 2D classification. An ab-initio structure was reconstructed and subjected to heterogeneous refinement (680,146 particles). Afterward, 675,529 particles were selected and subjected to NU refinement.

The particles were transferred to RELION and subjected to non-align local-masked 3D classification, using a focus mask covering YheS and the L1-stalk. First, 520,241 particles with YheS were selected, followed by further non-align local-masked 3D classification using a mask covering the A- and P-tRNAs. Then, 390,797 particles with the tRNAs were selected, followed by 3D refinement, CTF refinement, and Bayesian polishing. The particles were reprocessed with cryoSPARC and further subjected to non-align masked 3D classification, using a mask covering SSU and P-tRNA, resulting in the 3D classes with different SSU orientations against LSU. One class with 131,767 particles was selected. The codon-anticodon density of this class agrees with the expected SecM-stalled state, which has Pro and Gly tRNAs in the A and P sites, respectively.

To improve the occupancy of the factors, the particles were further subjected to non-align local-masked 3D classification, covering the A-tRNA. Afterward, 64,373 particles with A-tRNA were further subjected to non-align local-masked 3D classification, using a mask covering the A-tRNA, P-tRNA, YheS, and L1-stalk. After 50,797 particles were selected, they were subjected to NU refinement, resulting in the final map with 2.82 Å resolution.

To improve the local resolution, local-masked 3D refinements were performed using masks covering the SSU head, SSU body, LSU, and YheS with L1-stalk. The NU refined map and the local refined maps were subjected to B-factor sharpening and local resolution filtering. These maps were combined by Chimera (**55**) and used for model building and refinement.

### Model building and refinement

The reported *E. coli* ribosome structures, PDB IDs: 8QOA (**27**) and 8B0X (**45**), were used for the starting models of the RNA and protein parts, respectively. The YheS model was obtained by AlphaFold2 (**56**). The mRNA, tRNA, ATP, and nascent chain models were built manually. Manual revision was done using *Coot* 0.8 (**57**). Alternative conformations supported by the density were introduced. Geometrical restraints of modified residues and ligands were calculated by Grade Web Server (http://grade.globalphasing.org). The model was then refined against the composite map using Phenix.real_space_refine v1.20 (**58**) by global energy minimization and ADP refinement with rotamer and Ramachandran restraints.

Validation was performed by MolProbity (**59**). The statistics are listed in **Table S1**. ChimeraX (**60**) was used for making figures.

### β-galactosidase assay

*E. coli* cells harboring both pCY2524 (GFP-*secM*-*lacZ* reporter) and the YheS-expressing plasmid were grown overnight at 37°C in LB medium, supplemented with 100 µg/ml ampicillin and 20 µg/ml chloramphenicol. On the next day, they were inoculated into fresh LB medium containing 0.002% arabinose, 100 µg/ml ampicillin, and 20 µg/ml chloramphenicol and grown at 37°C. Expression of YheS or its derivatives was induced by the addition of 100 µM IPTG when the A_660_ reached 0.2. After further incubation (A_660_ = ∼0.6), 20 µl portions were subjected to a β-galactosidase assay as described previously (**10, 40**).

### Molecular simulation

The cryo-EM structure of the SecM-stalled ribosome was obtained from the Protein Data Bank (PDB: 8QOA) (**27**). To investigate the dynamics of SecM extension toward the C-terminus, we extracted the LSU RNAs within 10 Å of SecM (residues 149-165) and considered their connections, leaving a total of nine RNA strands (residues 745-752, 783-792, 1779-1783, 2055-2071, 2251-2253, 2438-2453, 2500-2507, 2572-2588, 2599-2612). This structure of SecM with the surrounding RNA was solvated, and counterions were added to maintain electroneutrality. The resulting system was subjected to 300-step energy minimization followed by 500-ps NVT and 500-ps NPT equilibration with heavy atom restraints. The force field parameters used were Amber ff19SB for proteins/peptides, TIP3P for water molecules, OL3 for RNA, and modrna08 for modified nucleosides. The equilibrated system was subjected to five 100-ns steered molecular dynamics (SMD) simulations, in which the distance between Lys149 (C_α_) on the N-terminus of SecM and Gly165 (C_α_) on the C-terminus was extended by 2, 5, or 10 Å, with the RNA backbone restrained. In addition, conventional MD simulations with only the C-terminus restrained were performed for another 200 ns; i.e., 100–300 ns. All molecular dynamics simulations were performed using the Amber 24 package (**61**).

## Funding

Platform Project for Supporting Drug Discovery and Life Science Research (Basis for Supporting Innovative Drug Discovery and Life Science Research (BINDS)) from AMED grant number JP25am121002, support no. 3272 (ON)

Japan Society for the Promotion of Science (JSPS) KAKENHI, Grant Number 22H02553 (YI)

Japan Society for the Promotion of Science (JSPS) KAKENHI, Grant Number 23H02410 (YC)

MEXT Grant-in-Aid for Scientific Research, Grant Number JP20H05925 (HT)

Japan Foundation for Applied Enzymology (YC)

Takeda Science Foundation (YC)

Yamada Science Foundation (YC)

Inamori Foundation (YC)

Noda Institute for Scientific Research (YC)

Senri Life Science Foundation (YC)

JST, PRESTO, Grant Number JPMJPR24OC (YC)

Core Research for Evolutional Science and Technology (CREST) of the Japan Science and Technology Agency (JST), Grant Number JPMJCR20E2 (ON).

## Author contributions

Each author’s contribution(s) to the paper should be listed (we suggest following the CRediT model with each CRediT role given its own line. No punctuation in the initials.

Examples:

Conceptualization: YC, YI

Methodology: TF, YC, YI

Investigation (cryo-EM): KI, YA, FKS, YI

Investigation (mutational analysis): KY, YC

Investigation (molecular dynamics): TI, TF

Visualization: KI, TI, YC

Supervision: HT, ON, YC, YI

Writing—original draft: YC, YI

Writing—review & editing: KI, TI, TF, HT, ON, YC, YI

## Data and materials availability

The atomic coordinates have been deposited in the Protein Data Bank (PDB) under the accession code 9VVI. The cryo-EM density maps have been deposited in the Electron Microscopy Data Bank (EMDB) under the accession code EMD-65381.

