## Supplementary Figures and Tables for "Structural insights into YheS-mediated release of SecM-arrested ribosome"

Kaishi Iso *et al.*

**This PDF file includes:**

Figs. S1 to S5
Tables S1 to S5

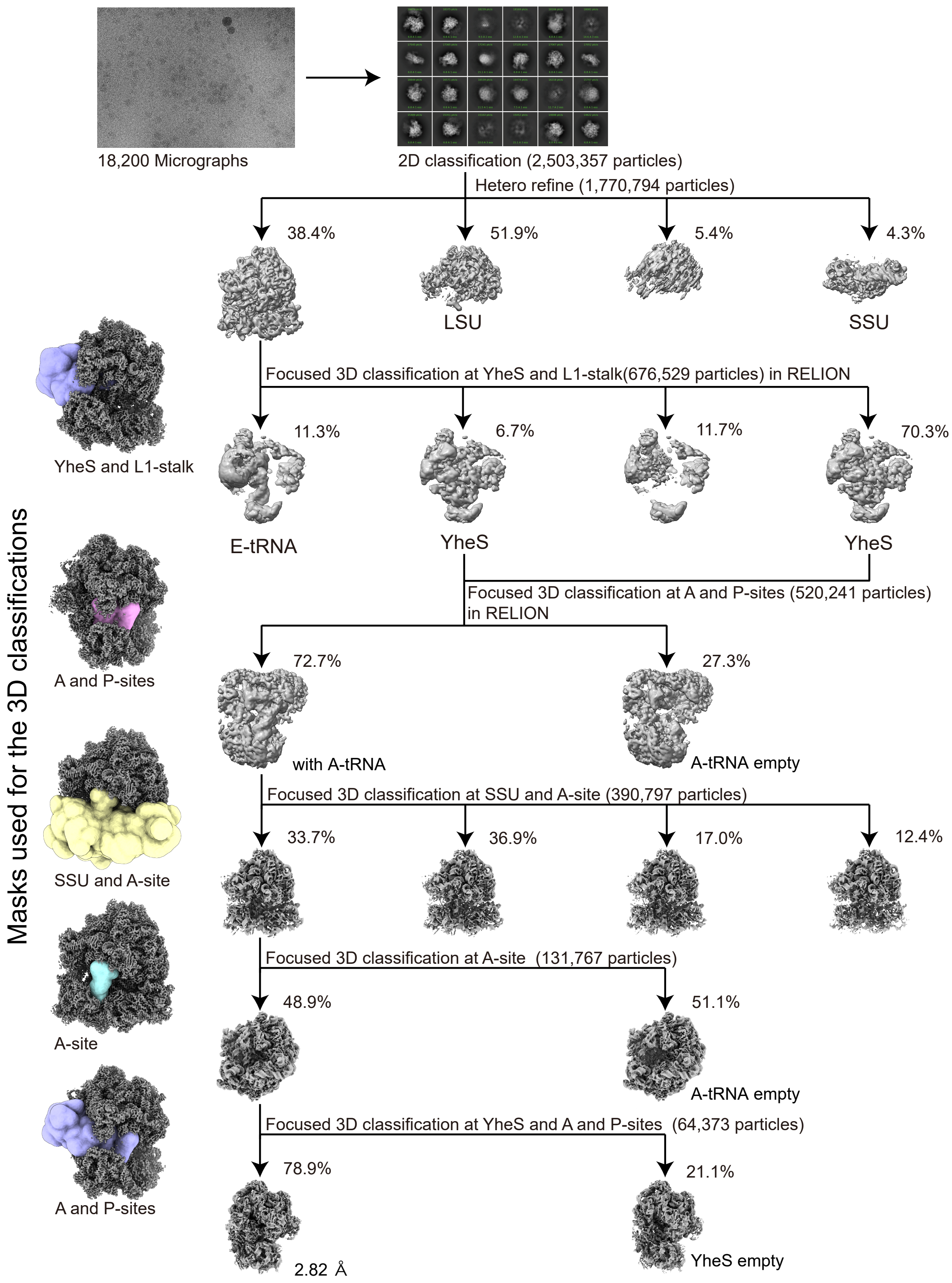

**Fig. S1. Cryo-EM data processing workflow.**

Schematic overview of the cryo-EM data processing pipeline. Local masks used for 3D classification without alignment are shown. See Methods for details.

### Overall (NU refine)

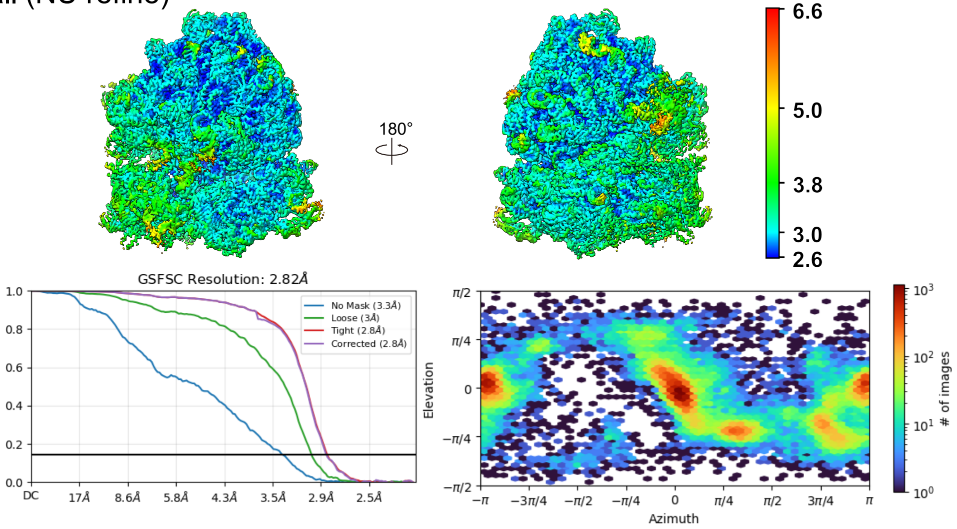

### SSU head (local-masked refined)

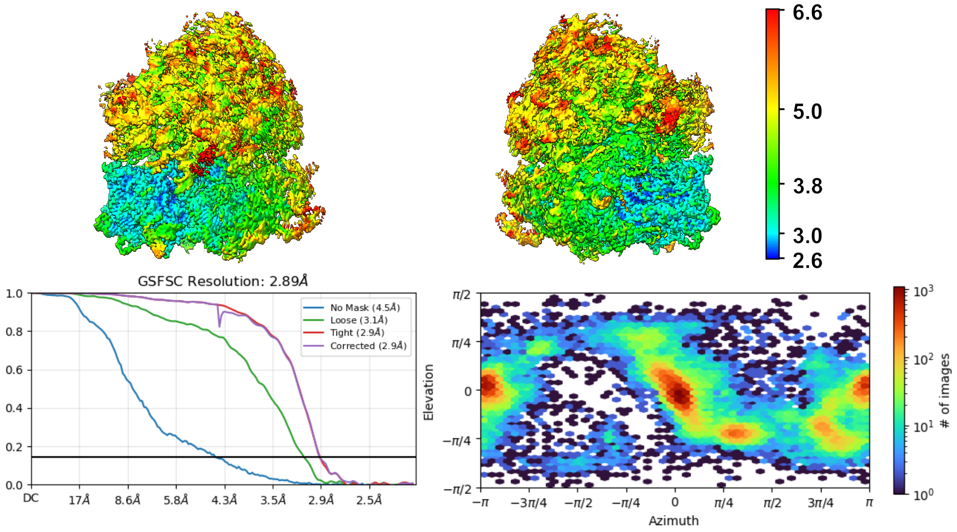

### SSU body (local-masked refined)

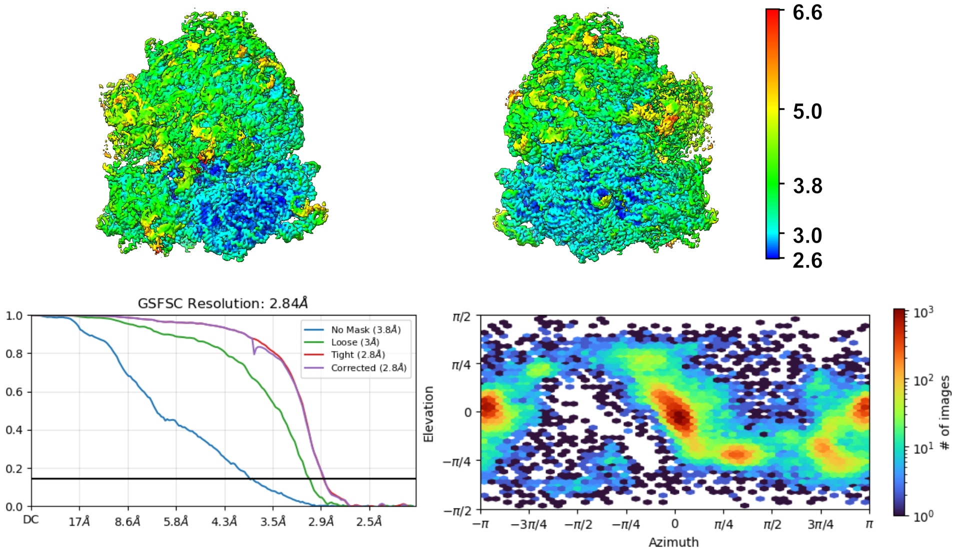

LSU (local-masked refined)

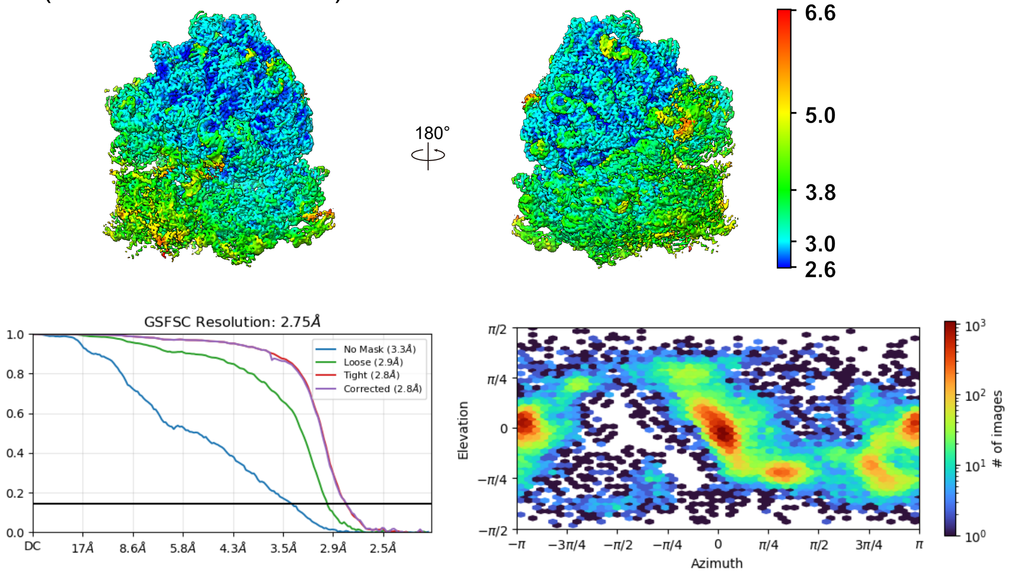

YheS and L1-stalk (local-masked refined)

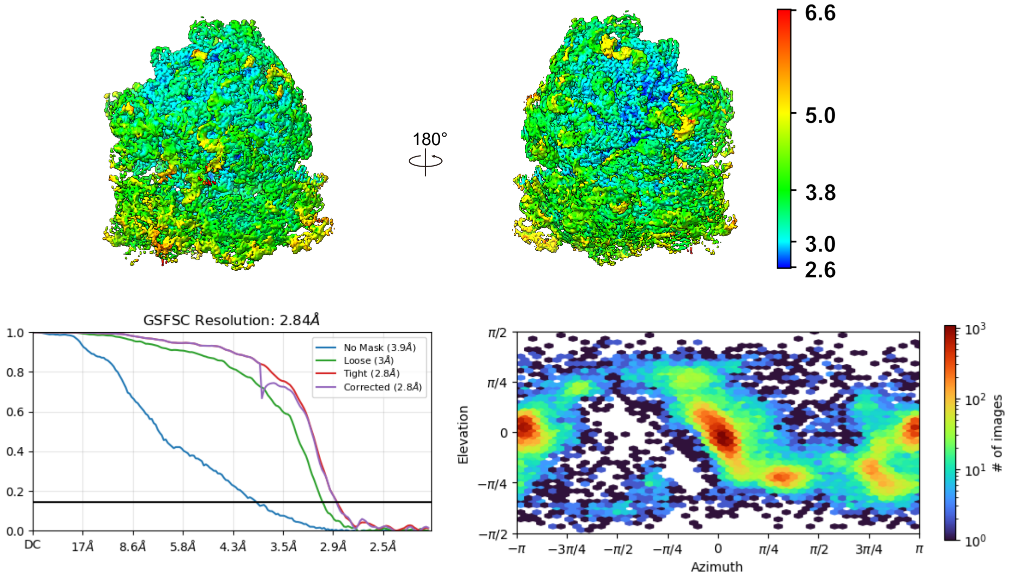

**Fig. S2. Quality of the cryo-EM reconstitutions.**

Cryo-EM maps from the overall non-uniform (NU) refinement and local-masked refinements, colored according to local resolution. Fourier shell correlation (FSC) curves of the half maps and the orientation distribution plot of the particles for each 3D reconstruction are shown.

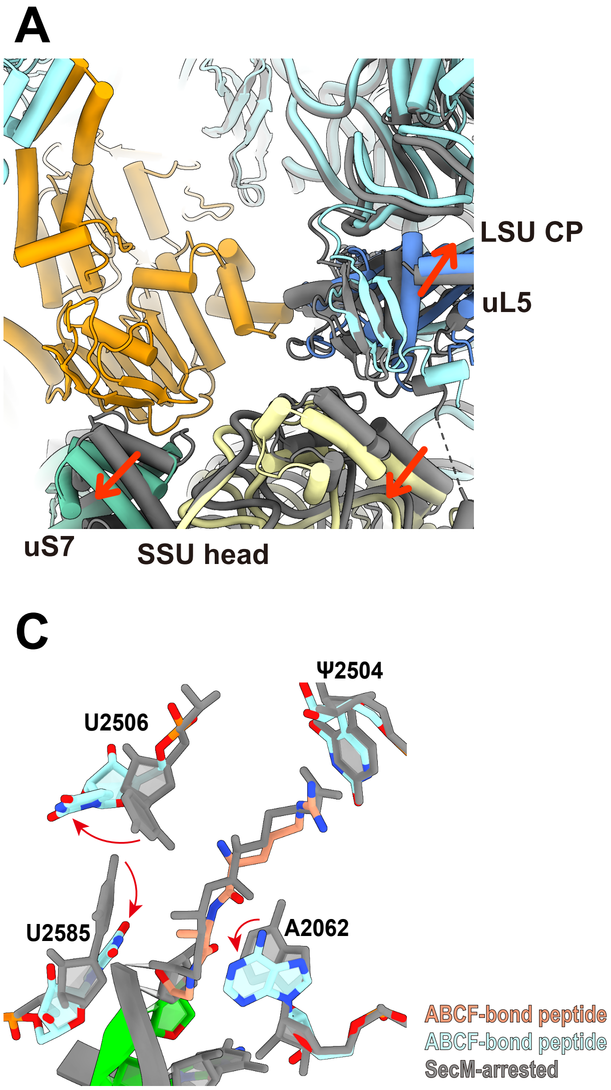

**Fig. S3. Structural rearrangement induced by YheS binding.**

(**A**) The SSU head undergoes a positional shift upon YheS binding, driven by its interaction with uS7, compared to the SecM-arrested ribosome without YheS (colored gray, PDB: 8QOA). The central protuberance (CP) of LSU is also shifted. These movements expand the E-site, where YheS engages the ribosome.

(**B**) The elbow of the P-tRNA is shifted by YheS, eliminating the contact between the P-tRNA elbow and uL5 in CP, which is observed in the SecM-arrested ribosome without YheS (colored gray, PDB: 8QOA).

(**C**) Rearrangement of the rRNA residues lining the peptide exti tunnel, compared to the SecM-arrested ribosome without YheS (colored gray, PDB: 8QOA).

(**D**) Rearrangement around the 3′-terminus of the P-tRNA. The base pairing between C75 of the P-tRNA and G2251 of the 23S rRNA and is disrupted, and A2602 of the 23S rRNA adopts a flipped-out conformation.

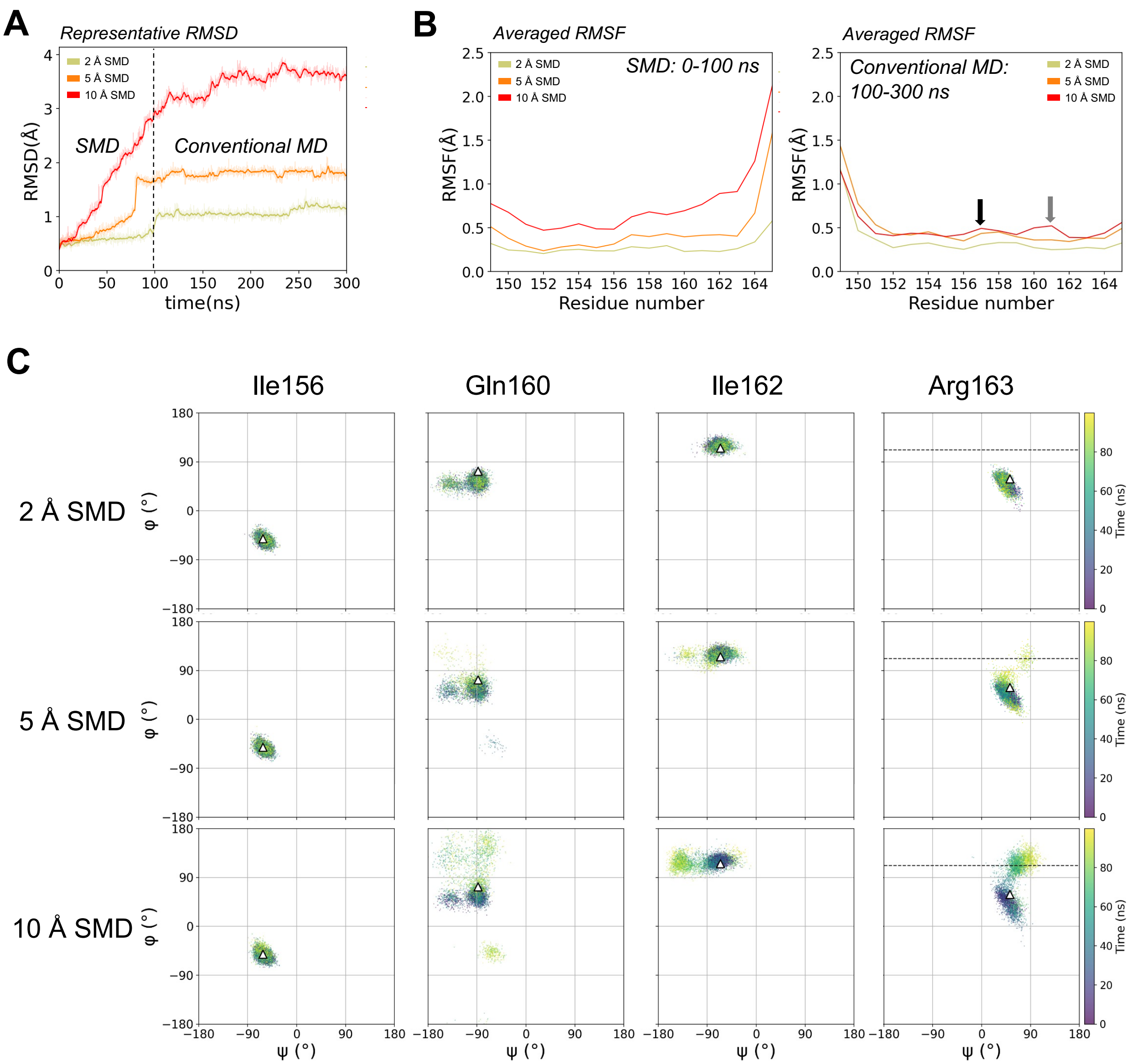

**Fig. S4. 100-ns SMD simulations of SecM and the subsequent 200-ns conventional MD simulations of SecM within the ribosome.**

(**A**) Representative root-mean-square deviation (RMSD) profiles of SecM during steered molecular dynamics (SMD) simulations (0–100 ns, under 2 Å, 5 Å, and 10 Å extensions) and subsequent conventional MD simulations (0–300 ns, only C-terminus restrained).
(**B**) Averaged root-mean-square fluctuation (RMSF) profiles of SecM from SMD simulation phase (0–100 ns, left) and the subsequent conventional MD simulation phase (100–300 ns, right). Black and gray arrows indicate slight increases in the structural fluctuations of S157 and Gly161, respectively.
(**C**) Changes in the backbone dihedral angles of four residues (Ile156, Gln160, Ile162, and Arg163) in five independent SMD simulations under the 2 Å, 5 Å and10 Å extensions. Each triangle indicates the position of the dihedral angles in the initial structure, and the dashed line in Arg163 represents the dihedral angle observed in the YheS-bound structure after arrest release.

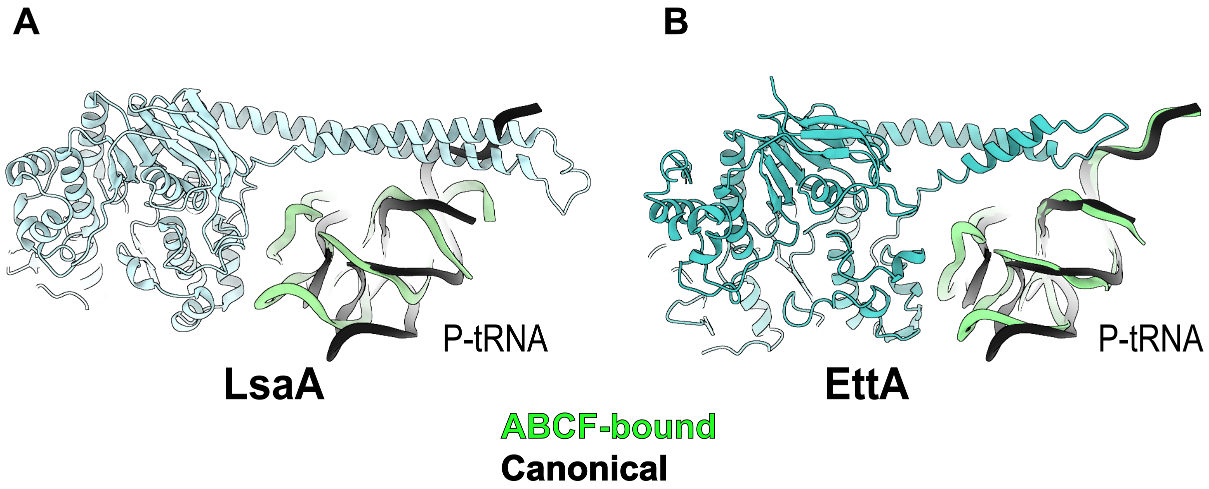

**Fig. S5. Comparison of the P-tRNA relocations induced by different ABCF proteins.** Structures of the ARE-ABCF LsaA (**A**, PDB: 7NHK, **reference-32**), and the non-ARE ABCF EttA (**B**, PDB: 7MSC, **reference-38**) are shown. In each panel, the P-tRNA from the ABCF-bound ribosome is colored green, while that from the canonical elongating ribosome (PDB: 8B0X) is shown in black for comparison. The P-tRNA relocation by EttA is smaller than that observed with LsaA.

**Table S1. Cryo-EM collection, processing, model refinement and validation statistics.**

| | |
|---|---|
| **Data collection and processing** | |
| Microscope | Titan Krios |
| Detector | K3 |
| Magnification | 165,000 |
| Voltage [kV] | 300 |
| Total electron exposure [$e^-$/Å$^2$] | 29.8 |
| Defocus range [μm] | −0.8 to −2.0 |
| Pixel size [Å] | 0.83 |
| Number of micrographs | 18,200 |
| Symmetry imposed | $C_1$ |
| Final particles | 50,797 |
| Resolution [Å] (Overall/LSU/SSU body/SSU head/YheS and L1 stalk) | 2.82/2.75/2.84/2.89/2.84 |
| Map-sharpening $B$ factor [Å$^2$] (Overall/LSU/ SSU body/SSU head/YheS and L1 stalk) | −56.6/−58.2/−58.5/−57.8/−55.4 |
| **Refinement** | |
| Model composition | |
| Total atoms | 150,960 |
| Chains (RNA/protein) | 6/52 |
| RNA residues (non-modified/modified) | 4,612/47 |
| Protein residues (non-modified/modified) | 6,444/5 |
| Metal ions ($Mg^{2+}$/$K^+$/$Zn^{2+}$) | 265/125/2 |
| Ligands (ATP/spermidine) | 2/3 |
| Waters | 18 |
| Model to map CC ($CC_{mask}$/$CC_{box}$/$CC_{peaks}$/$CC_{volume}$) | 0.89/0.91/0.86/0.89 |
| Resolution [Å] by model-to-map FSC, threshold 0.50 (masked/ unmasked) | 2.84/2.84 |
| Average $B$ factor [Å$^2$] (Overall/RNA/protein/metal ion and ligand/water) | 80.7/90.5/59.5/46.6 |
| R.m.s. deviations, bond lengths [Å]/bond angles [°] | 0.002/ 0.461 |
| **Validation** | |
| Clash score | 6.49 |
| Rotamer outliers [%] | 0.19 |
| Ramachandran plot [%] (Favored/allowed/disallowed) | 98.66/1.33/0.02 |
| CaBLAM outliers [%] | 1.09 |
| $C_\beta$ outliers [%] | 0 |
| EMRinger score | 4.14 |
| MolProbity score | 1.36 |

**Table S2. Polar and stacking interactions between YheS and ribosome.**

| YheS Region | Residue | Moiety | Ribosome region | Subunit/Domain | Residue | Moiety | Interaction type |
|---|---|---|---|---|---|---|---|
| Arm | Asn105 | Side chain | L1 stalk | uL1 | Arg122 | Side chain | π stacking |
| Arm | His114 | Side chain | L1 stalk | uL1 | Gly125 | Main chain | H bond |
| Arm | Asp118 | Side chain | L1 stalk | uL1 | Arg164 | Side chain | H bond/salt bridge |
| Arm | Asp121 | Side chain | L1 stalk | uL1 | Lys167 | Side chain | H bond/salt bridge |
| NBD1 | Trp123 | Side chain | L1 stalk | uL1 | Gly132 | Main chain | π stacking |
| NBD1 | Ser127 | Side chain | L1 stalk | 23S rRNA H76 | G2112 | Base | H bond |
| NBD1 | His134 | Side chain | L1 stalk | 23S rRNA H76 | U2112 | Ribose | H bond |
| NBD1 | Asn140 | Side chain | L1 stalk | 23S rRNA H76 | G2113 | Ribose | H bond |
| NBD1 | Asn25 | Side chain | 50S body | 23S rRNA H68 | U1886 | Phosphate | H bond |
| NBD1 | Thr228 | Side chain | 50S body | 23S rRNA H68 | U1851 | Phosphate | H bond |
| PtIM | Gln244 | Side chain | 50S body | 23S rRNA H68 | C1893 | Ribose | H bond |
| NBD2 | Asn310 | Main chain | 30S head | uS7 | Gly112 | Main chain | H bond |
| NBD2 | Asn310 | Side chain | 30S head | uS7 | Lys110 | Main chain | H bond |
| NBD2 | Asn334 | Side chain | 30S head | uS7 | Gly132 | Main chain | H bond |
| NBD2 | Ser339 | Side chain | 30S head | uS7 | Lys136 | Side chain | H bond |
| NBD2 | Asp496 | Side chain | 30S head | uS7 | Arg143 | Side chain | H bond/salt bridge |
| PtIM | His258 | Side chain | 50S body | 23S rRNA H93 | U2596 | Phosphate | H bond |
| PtIM | Arg267 | Side chain | 50S body | 23S rRNA H80 | G2255 | Phosphate | H bond/salt bridge |
| PtIM | Thr271 | Side chain | 50S body | 23S rRNA H93 | A2602 | Phosphate | H bond |
| PtIM | Lys274 | Side chain | 50S body | 23S rRNA H74 | G2437 | Phosphate | H bond/salt bridge |
| PtIM | Gln275 | Side chain | 50S body | 23S rRNA H74 | G2436 | Ribose | H bond |
| PtIM | Gln275 | Side chain | 50S body | 23S rRNA H93 | G2595 | Ribose | H bond |
| PtIM | Gln275 | Side chain | 50S body | 23S rRNA H93 | G2597 | Base | H bond |
| PtIM | Gln277 | Side chain | 50S body | 23S rRNA H80 | G2255 | Phosphate | H bond |
| PtIM | Ser278 | Side chain | 50S body | 23S rRNA H74 | G2435 | Ribose | H bond |
| PtIM | Arg279 | Side chain | 50S body | 23S rRNA H93 | U2595 | Ribose | H bond |
| PtIM | Arg279 | Side chain | 50S body | 23S rRNA H93 | G2597 | Base | H bond |
| PtIM | Lys281 | Side chain | 50S body | 23S rRNA H74 | G2436 | Phosphate | H bond/salt bridge |
| PtIM | Gln285 | Side chain | 50S body | 23S rRNA H75 | A2077 | Ribose | H bond |

**Table S3. Polar and stacking interactions between YheS and P-tRNA.**

| YheS region | Residue | Moiety | P-tRNA region | Residue | Moiety | Interaction type |
|---|---|---|---|---|---|---|
| PtIM | Lys269 | Side chain | Acceptor arm | U73 | Phosphate | H bond |
| PtIM | Ala270 | Main chain | Acceptor arm | C74 | Ribose | H bond |
| NBD2 | Gln408 | Side chain | Elbow | C56 | Ribose | H bond |
| NBD2 | Gln408 | Side chain | Elbow | C57 | Ribose | H bond |
| NBD2 | Arg411 | Side chain | Elbow | C56 | Base | π stacking |
| NBD2 | Asp412 | Side chain | Elbow | G19 | Base | H bond |
| NBD2 | Gly421 | Main chain | Elbow | G19 | Base | H bond |

**Table S4. Used Plasmids.**

| plasmid name | feature | source | Construction_Quick change PCR / Gibson assembly | | | Construction_Gibson assembly_fragment_2 | | |
|---|---|---|---|---|---|---|---|---|
| | | | primer 1 | primer 2 | template DNA-1 | primer 1 | primer 2 | template DNA |
| pBAD30 | $Amp^R$, p15A ori, $P_{BAD}$ expression system | Guzman et al., 1995 | | | | | | |
| pCA24N | $Cm^R$, ColE1 ori, lacO - $P_{T5}$ expression system | Kitagawa et al., 2005 | | | | | | |
| pCY1061 | pCA24N carrying His6-sfGFP-secM-lacZα | Chadani et al., 2017 | | | | | | |
| pCY2524 | pBAD30 carrying sfGFP-secM(E134-R168)-lacZ | Chadani et al., 2024 | | | | | | |
| pCY3863 | pBAD30 carrying His6-yheS (E175Q, E456Q) | Chadani et al., 2024 | | | | | | |
| pCY3867 | pCA24N carrying folA_60aa-secM(E134-R168)-lacZa | This study | PT2061 | PM1043 | *E. coli* genome | PM0292 | PT0123 | pCY1061 |
| pCY3893 | pCA24N carrying His6-yheS | Chadani et al., 2024 | | | | | | |
| pCY4476 | derivative of pCY3893 carrying His6-*yheS* (N25A) | This study | PT3050 | PT3051 | pCY3893 | | | |
| pCY4481 | derivative of pCY3893 carrying His6-*yheS* (D496A) | This study | PT3058 | PT3059 | pCY3893 | | | |
| pKY0003 | derivative of pCY3893 carrying His6-*yheS* (K269A) | This study | KY0005 | KY0006 | pCY3893 | | | |
| pKY0009 | derivative of pCY3893 carrying His6-*yheS* (Q275A) | This study | KY0017 | KY0018 | pCY3893 | | | |
| pKY0011 | derivative of pCY3893 carrying His6-*yheS* (Q277A) | This study | KY0021 | KY0022 | pCY3893 | | | |
| pKY0013 | derivative of pCY3893 carrying His6-*yheS* (R279A) | This study | KY0025 | KY0026 | pCY3893 | | | |
| pKY0015 | derivative of pCY3893 carrying His6-*yheS* (K281A) | This study | KY0029 | KY0030 | pCY3893 | | | |
| pKY0044 | derivative of pCY3893 carrying His6-*yheS* (N310A) | This study | KY0089 | KY0090 | pCY3893 | | | |
| pKY0091 | derivative of pCY3893 carrying His6-*yheS* (T228A) | This study | KY0177 | KY0178 | pCY3893 | | | |
| pKY0107 | derivative of pCY3893 carrying His6-*yheS* (Q244A) | This study | KY0209 | KY0210 | pCY3893 | | | |
| pKY0228 | derivative of pCY3893 carrying His6-*yheS* (Q408A) | This study | KY0334 | KY0335 | pCY3893 | | | |
| pKY0229 | derivative of pCY3893 carrying His6-*yheS* (R411A) | This study | KY0336 | KY0337 | pCY3893 | | | |
| pKY0230 | derivative of pCY3893 carrying His6-*yheS* (D412A) | This study | KY0338 | KY0339 | pCY3893 | | | |
| pKY0231 | derivative of pCY3893 carrying His6-*yheS* (Q408A R411A D412A) | This study | KY0335 | KY0340 | pCY3893 | | | |
| pKY0232 | derivative of pCY3893 carrying His6-*yheS* (A270P) | This study | KY0341 | KY0008 | pCY3893 | | | |
| pKY0233 | derivative of pCY3893 carrying His6-*yheS* (K269A A270P) | This study | KY0342 | KY0006 | pCY3893 | | | |

| | | | | | |
|---|---|---|---|---|---|
| pKY0234 | derivative of pCY3893 carrying His6-*yheS* (S127A) | This study | KY0343 | KY0344 | pCY3893 |
| pKY0235 | derivative of pCY3893 carrying His6-*yheS* (H134A) | This study | KY0345 | KY0346 | pCY3893 |
| pKY0236 | derivative of pCY3893 carrying His6-*yheS* (N140A) | This study | KY0347 | KY0348 | pCY3893 |
| pKY0237 | derivative of pCY3893 carrying His6-*yheS* (S127A H134A N140A) | This study | KY0349 | KY0350 | pCY3893 |
| pKY0238 | derivative of pCY3893 carrying His6-*yheS* (H114A) | This study | KY0351 | KY0352 | pCY3893 |
| pKY0239 | derivative of pCY3893 carrying His6-*yheS* (D118A) | This study | KY0353 | KY0354 | pCY3893 |
| pKY0240 | derivative of pCY3893 carrying His6-*yheS* (W123A) | This study | KY0355 | KY0356 | pCY3893 |
| pKY0241 | derivative of pCY3893 carrying His6-*yheS* (H114A D118A W123A) | This study | KY0357 | KY0358 | pCY3893 |
| pKY0242 | derivative of pCY3893 carrying His6-*yheS* (Q275A Q277A R279A K281A) | This study | KY0359 | KY0018 | pCY3893 |
| pKY0308 | derivative of pCY3893 carrying His6-*yheS* (N334A) | This study | PT3074 | PT3075 | pCY3893 |
| pKY0309 | derivative of pCY3893 carrying His6-*yheS* (S339A) | This study | PT3076 | PT3077 | pCY3893 |
| pKY0310 | derivative of pCY3893 carrying His6-*yheS* (N310A D496A) | This study | KY0089 | KY0090 | pCY4481 |
| pKY0311 | derivative of pCY3893 carrying His6-*yheS* (N310A N334A S339A D496A) | This study | PT3072 | PT3073 | pKY0310 |

**Table S5. Used oligonucleotides.**

| Oligonucleotide name | sequence (5' to 3') |
|---|---|
| **PM0292** | CTGGTTCCGCGTGGATCCACGAATTCGAGCTCGGTA |
| **PM1043** | GGATCCACGCGGAACCAGAATATTTTTGCGTCCTGGCAAC |
| **PT0123** | CATAGTTAATTTCTCCTCTTTAATGAATTC |
| **PT2061** | GAGGAGAAATTAACTATGAAGTACTATATCAGTCTGATTGCGGCG |
| **PT3050** | AATGCCACCGCCACCATCGCACCTGGGCAGAAAGTCGGC |
| **PT3051** | GATGGTGGCGGTGGCATTATCCAGCAGGACGCGCAC |
| **PT3058** | CTGCGTTCCACCACTGACGCTCTCTACCTGGTTCACGA |
| **PT3059** | GTCAGTGGTGGAACGCAGCAAATGACGGTCGTGCGA |
| **PT3072** | GCCCTGGTGCCCGGCGCACGTATTGGTCTGTTAGGC |
| **PT3073** | TGCGCCGGGCACCAGGGCCAGTTTAATCGAGTCGAG |
| **PT3074** | CTCGACTCGATTAAACTGGCCCTGGTGCCCGGCTCGCGT |
| **PT3075** | CAGTTTAATCGAGTCGAGAATAATGCGATCGCCATAG |
| **PT3076** | CTGAACCTGGTGCCCGGCGCACGTATTGGTCTGTTAGGC |
| **PT3077** | GCCGGGCACCAGGTTCAGTTTAATCGAGTCGAGAATAATG |
| **KY0005** | ATCGACCGTTTCCGTGCCGCAGCCACCAAAGCGAAGCAG |
| **KY0006** | GGCACGGAAACGGTCGATATAACTTTGCAGATGCGCTAC |
| **KY0008** | TTTGGCACGGAAACGGTCGATATAACTTTGCAGATGC |
| **KY0017** | AAAGCCACCAAAGCGAAGGCGGCCCAGAGCCGCATTAAG |
| **KY0018** | CTTCGCTTTGGTGGCTTTGGCACGGAAACGGTCGATATAAC |
| **KY0021** | ACCAAAGCGAAGCAGGCCGCGAGCCGCATTAAGATGCTCGAGCG |
| **KY0022** | GGCCTGCTTCGCTTTGGTGGCTTTGGCACGGAAACG |
| **KY0025** | GCGAAGCAGGCCCAGAGCGCCATTAAGATGCTCGAGCGTATG |
| **KY0026** | GCTCTGGGCCTGCTTCGCTTTGGTGGCTTTGGCACG |
| **KY0029** | CAGGCCCAGAGCCGCATTGCGATGCTCGAGCGTATGGAG |
| **KY0030** | AATGCGGCTCTGGGCCTGCTTCGCTTTGGTGGCTTTG |
| **KY0089** | GCGCCGGAAAGCCTGCCAGCTCCGTTACTGAAGATGGAAAAAG |
| **KY0090** | TGGCAGGCTTTCCGGCGCGCGGAAGCTAAAGCGGAAC |
| **KY0177** | CAAAGCATGTTCGAGTACGCCGGCAACTACAGTTCGTTTGAAG |
| **KY0178** | GTACTCGAACATGCTTTGTTGTTCGATATGAATAATTTTATCGAC |

| | |
|---|---|
| **KY0209** | CGCGCCACCCGTCTGGCGGCGCAACAAGCGATGTACGAAAG |
| **KY0210** | CGCCAGACGGGTGGCGCGCTGTACTTCAAACGAAC |
| **KY0334** | GCGCCGCAGGAGCTGGAAGCAAAACTGCGTGACTACCTC |
| **KY0335** | TTCCAGCTCCTGCGGCGCTAAACGTGCCAGATGTTG |
| **KY0336** | GAGCTGGAACAAAAACTGGCTGACTACCTCGGCGGCTTTGG |
| **KY0337** | CAGTTTTTGTTCCAGCTCCTGCGGCGCTAAACGTGC |
| **KY0338** | CTGGAACAAAAACTGCGTGCCTACCTCGGCGGCTTTGGTTTC |
| **KY0339** | ACGCAGTTTTTGTTCCAGCTCCTGCGGCGCTAAACG |
| **KY0340** | GCGCCGCAGGAGCTGGAAGCAAAACTGGCTGCCTACCTCGGCGGCTTTGGTTTC |
| **KY0341** | GACCGTTTCCGTGCCAAACCCACCAAAGCGAAGCAGGCCC |
| **KY0342** | ATCGACCGTTTCCGTGCCGCACCCACCAAAGCGAAGCAGGCCC |
| **KY0343** | GACGCATGGAGTATTCGCGCCCGTGCTGCCAGCCTGCTGC |
| **KY0344** | GCGAATACTCCATGCGTCAATAGCATCCAGCTTGCC |
| **KY0345** | CGTGCTGCCAGCCTGCTGGCCGGCCTCGGTTTCAGCAATG |
| **KY0346** | CAGCAGGCTGGCAGCACGGGAGCGAATACTCCATGC |
| **KY0347** | CACGGCCTCGGTTTCAGCGCTGAACAACTGGAGCGCCCG |
| **KY0348** | GCTGAAACCGAGGCCGTGCAGCAGGCTGGCAGCA |
| **KY0349** | AGCCTGCTGGCCGGCCTCGGTTTCAGCGCTGAACAACTGGAGCGCCCG |
| **KY0350** | CCGAGGCCGGCCAGCAGGCTGGCAGCACGGGCGCGAATACTCCATGCGTC |
| **KY0351** | CACGCCATTGCGACCATTGCTGGCAAGCTGGATGCTATTG |
| **KY0352** | AATGGTCGCAATGGCGTGCCCGTCGTTACGTTCGTTG |
| **KY0353** | ACCATTCATGGCAAGCTGGCTGCTATTGACGCATGGAGTATTC |
| **KY0354** | CAGCTTGCCATGAATGGTCGCAATGGCGTGCCCGTC |
| **KY0355** | CTGGATGCTATTGACGCAGCGAGTATTCGCTCCCGTGCTGCC |
| **KY0356** | TGCGTCAATAGCATCCAGCTTGCCATGAATGGTCGC |
| **KY0357** | CAAGCTGGCTGCTATTGACGCAGCGAGTATTCGCTCCCGTGCTGCC |
| **KY0358** | CGTCAATAGCAGCCAGCTTGCCAGCAATGGTCGCAATGGCGTG |
| **KY0359** | AAAGCCACCAAAGCGAAGGCGGCCGCGAGCGCCATTGCGATGCTCGAGCGTATG GAG |
